# Comparison of concurrent and asynchronous running kinematics and kinetics from marker-based motion capture and markerless motion capture under two clothing conditions

**DOI:** 10.1101/2023.02.22.529537

**Authors:** Robert M. Kanko, Jereme B. Outerleys, Elise K. Laende, W. Scott Selbie, Kevin J. Deluzio

**Author notes:** Declaration of Interests: RMK and JBO are employed by Theia Markerless Inc, and WSS is the CEO of Theia Markerless Inc. Data for this study was collected by RMK in the Human Mobility Research Laboratory at Queen’s University under the supervision of KJD, prior to employment with Theia Markerless Inc.

## Abstract

As markerless motion capture is increasingly used to measure three-dimensional human pose, it is important to understand how markerless results can be interpreted alongside historical marker-based data and how they are impacted by clothing. We compared concurrent running kinematics and kinetics between marker-based and markerless motion capture, and between two markerless clothing conditions. Thirty adults ran on an instrumented treadmill wearing motion capture clothing while concurrent marker-based and markerless data were recorded, and ran a second time wearing athletic clothing (shorts and t-shirt) while markerless data were recorded. Differences calculated between the concurrent signals from both systems, and also between each participant’s mean signals from both asynchronous clothing conditions were summarized across all participants using root-mean-square differences. Most kinematic and kinetic signals were visually consistent between systems and markerless clothing conditions. Between systems, joint center positions differed by 3 cm or less, sagittal plane joint angles differed by 5° or less, and frontal and transverse plane angles differed by 5°-10°. Joint moments differed by 0.3 Nm/kg or less between systems. Differences were sensitive to segment coordinate system definitions, highlighting the effects of these definitions when comparing against historical data or other motion capture modalities.

## 1. Introduction

Motion capture has long been used to quantify movement patterns, monitor interventions and has potential as a tool for injury prediction. The latter has been a particular focus for those interested in its application to running biomechanics. However, due to the inherent time, cost, environment, and expertise constraints associated with traditional marker-based motion capture approaches, the successes of this tool have been limited. Likewise, wearable sensors such as inertial measurement units (IMUs) have limitations, specifically related to magnetic field effects such as those generated by laboratory treadmills^1,2^ and are best suited to reporting quantities such as accelerations and angular rates which they measure directly, rather than estimating intersegmental measures such as joint angles.

As emerging technology, markerless motion capture has the potential to capture high resolution three dimension (3D) biomechanical signals of running in a more accessible system that avoids potential effects of applying markers or sensors. Ongoing developments, validations, and applications demonstrate the promise of this form of motion capture technology for increasing the usefulness of motion capture-based running gait analysis. Markerless motion capture systems such as Theia3D (Theia Markerless Inc., Kingston, Ontario, Canada) can measure kinematics using multiple synchronized video cameras with few restrictions on the data collection environment, subject attire, and a significantly reduced collection time compared to marker-based motion capture. Markerless systems have demonstrated success in measuring comparable kinematic signals to those from marker-based motion capture during walking^3,4^, boxing^5^, and running^6^. While marker-based motion capture may not provide a true gold standard for biomechanical data collection due to issues with soft tissue artefact and marker placement variation, it is a historically accepted technology and forms the basis of much of the running biomechanics literature. Understanding how data collected with markerless motion capture compares to marker-based data, as well as the factors contributing to any differences between the two systems, is an important step in the introduction of markerless technology to the field of running biomechanics. Further, we were motivated to understand the effect of clothing on the reliability of the markerless system. Therefore, the objectives of this study were to compare kinematic and kinetic data from concurrent markerless and marker-based motion capture during treadmill running, examine the impact of clothing on markerless motion capture, and investigate the sensitivity of kinematic and kinetic data to biomechanical model definitions.

## 2. Methods

### 2.1 Data Collection

Thirty healthy, active young adults (15f/15m, mean (SD) age: 23.0 (3.5) years, height: 1.76 (0.09) m, mass: 69.2 (11.4) kg, BMI: 22.3 (3.0) kg/m^2^) were recruited from the university community to participate in this study at the Human Mobility Research Laboratory (Kingston, Ontario, Canada) (Figure 1). Exclusion criteria included having any lower-limb musculoskeletal injury in the past six months or any neuromuscular or musculoskeletal impairments that could hinder their performance of running. The research protocol was approved by the institutional research ethics board, and participants provided written informed consent.

**Figure 1:**
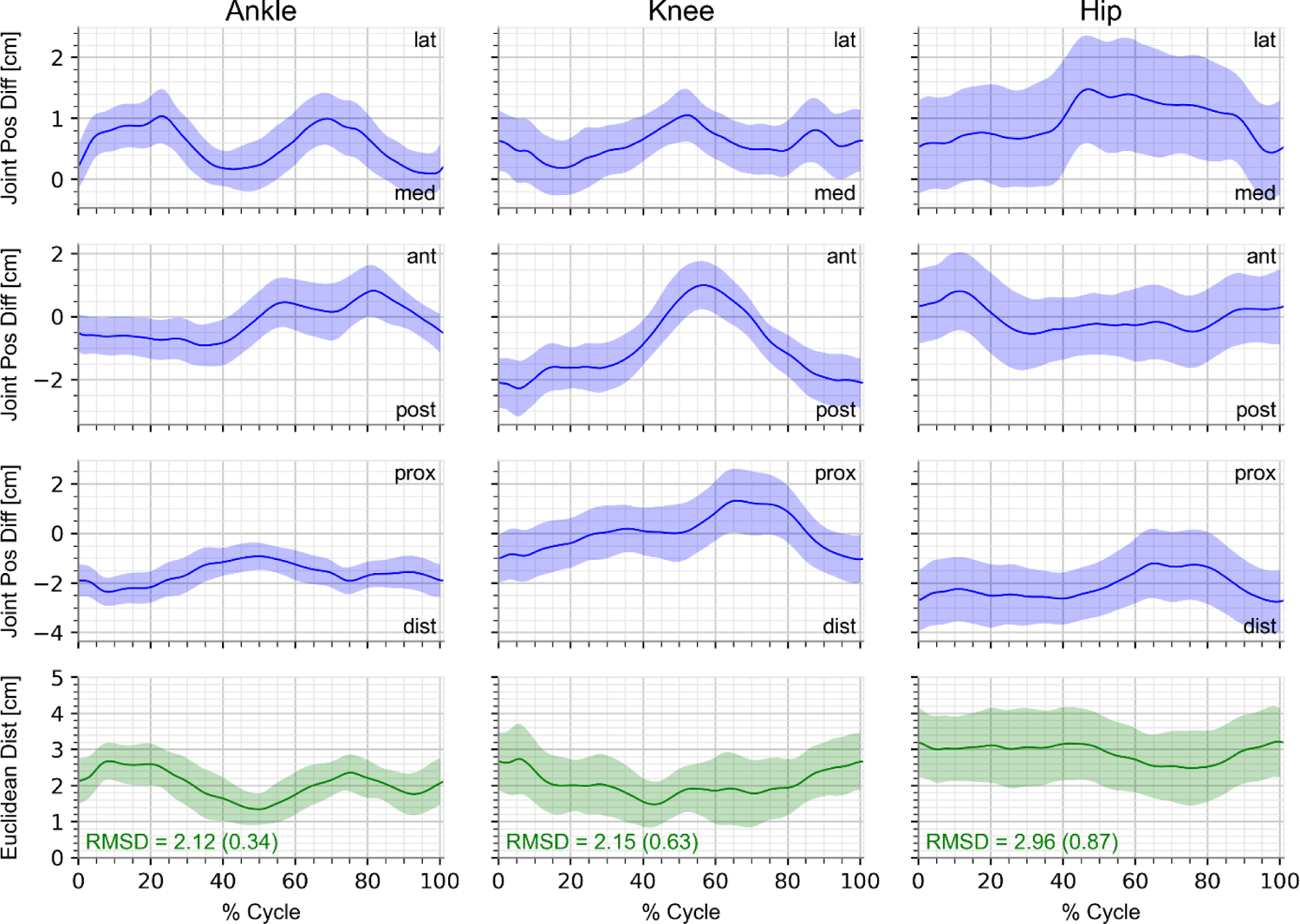
Mean joint center position differences between concurrent marker-based motion capture and markerless MoCap clothing condition across all participants, for the global x-direction (row 1; lateral/medial), global y-direction (row 2; anterior/posterior), global z-direction (row 3; proximal/distal), and 3D Euclidean distances (row 4). Differences were assessed as markerless joint position - marker-based joint position; inset position descriptors are for the markerless joint center relative to the marker-based joint center. Mean (SD) RMS of the 3D Euclidean distances is inset for each joint.

A marker-based motion capture system consisting of seven Qualisys Oqus marker-based cameras (Qualisys AB, Gothenburg, Sweden) and a markerless motion capture system of eight Qualisys Miqus video cameras (full HD 1920×1080 resolution, RBG video) were arranged around an instrumented treadmill (AMTI Inc, Watertown, MA) and were connected to a single instance of Qualisys Track Manager (QTM), ensuring their temporal synchronization^7^. Both sets of cameras were calibrated and localized simultaneously (calibration residual errors < 1 mm), ensuring they shared one global coordinate system (y-axis pointing in the treadmill direction of progression, x-axis pointing right laterally from the direction of progression, and z-axis pointing vertically). Both camera systems recorded at 85 Hz, the maximum frame rate for the Miqus cameras at Full HD (1920×1080) resolution. This camera resolution and frame rate combination was selected to maximize the resolution used to capture participants, which limited the frame rate to 85 Hz. Ground reaction forces were recorded at 425 Hz.

For the marker-based system, a static calibration trial was recorded of the subject standing on the treadmill with their feet pointing anteriorly and positioned approximately shoulder-width apart, their elbows flexed to approximately 90 degrees, and their palms facing the floor. The markerless motion capture system does not use a static calibration trial.

Participants ran at a self-selected speed under two clothing conditions: (1 - ‘MoCap clothing’) wearing typical marker-based motion capture clothing and markers^7^, and (2 - ‘Sport clothing’) wearing self-selected athletic clothing which typically consisted of a short-sleeved or sleeveless shirt and shorts that end at or above the knee. Half of the participants performed the MoCap condition first and half performed the Sport condition first; the order was not randomized across the participants. Participants selected their running speed by starting from stationary and providing feedback to the experimenters to increase or decrease the speed until they were satisfied with the current speed. The current speed was recorded by the experimenters and was replicated during the second condition without conveying its value to the participant at any time.

During the MoCap clothing condition, marker-based motion capture, markerless motion capture, and ground reaction force data were recorded simultaneously. During the Sport clothing condition, markerless motion capture and ground reaction force data were recorded. During both conditions, participants wore their personal running shoes and were allowed to acclimatize to their self-selected running speed on the treadmill for two minutes before one 10-second trial was recorded. Concurrent marker-based and markerless walking data under the MoCap clothing condition also obtained during these data collections have been analyzed and published previously^3,7^.

### 2.2 Data Analysis

Markerless video data were processed in Theia3D v2022-1-0-2309 (Theia Markerless Inc, Kingston, Ontario, Canada), using 3 degrees of freedom (DOF) at the hip, knee, and ankle joints, and a 20 Hz equivalent GCVSPL cut-off frequency^8^. Marker-based motion capture data were tracked in QTM and exported for further analysis in Visual3D (C-Motion Inc, Germantown, MD). Marker trajectories were filtered using a 20 Hz equivalent GCVSPL cut-off frequency, and a skeletal model was manually defined with similar segment coordinate system definitions to the Theia3D model. Model segment lengths were independent between systems, being based on the markers for the marker-based model and being automatically generated for the markerless model. The marker-based skeletal model used identical inverse kinematic segment chains and joint constraints to those used by Theia3D (Table 1).

**Table 1:** Marker-based model lower body segment coordinate system definitions and inverse kinematic model constraints, selected to best match the segment definitions and constraints of the Theia3D model.

| Segment | Coordinate System | Description |
| --- | --- | --- |
| Pelvis | Origin | Midpoint between mid-ASIS and mid-PSIS |
|  | X-Axis | Left ASIS to right ASIS |
|  | Y-Axis | Mid-PSIS to mid-ASIS |
|  | Z-Axis | Normal to xy-plane |
|  | Parent, DOF | Lab, 6 DOF |
| Kinematic (Virtual) Pelvis | Origin | Point at the height of midpoint between left and right iliac crest markers along vector between midpoint between left and right hip and midpoint between C7 and sternal notch markers |
|  | X-Axis | Left hip joint center to right hip joint center |
|  | Y-Axis | Normal to xz-plane |
|  | Z-Axis | Segment origin to midpoint between C7 and sternal notch markers |
|  | Parent, DOF | Pelvis, 0 DOF |
| Thigh | Origin | Hip joint center of Pelvis, based on (Bell et al., 1990) |
|  | X-Axis | Vector perpendicular to z-axis lying in the plane formed by hip joint center, medial and lateral femoral epicondyles |
|  | Y-Axis | Vector normal to plane formed by hip joint center, medial and lateral femoral epicondyles |
|  | Z-Axis | Midpoint between medial and lateral femoral epicondyles to hip joint center |
|  | Parent, DOF | Pelvis, 3 DOF |
| Shank | Origin | Midpoint between medial and lateral femoral epicondyles |
|  | X-Axis | Vector perpendicular to z-axis lying in the plane formed by segment origin, medial and lateral malleoli |
|  | Y-Axis | Vector normal to plane formed by segment origin, medial and lateral malleoli |
|  | Z-Axis | Midpoint between medial and lateral malleoli to segment origin |
|  | Parent, DOF | Thigh, 3 DOF |
| Foot | Origin | Midpoint between medial and lateral malleoli |
|  | X-Axis | Vector normal to y- and z-axes |
|  | Y-Axis | Segment origin to segment distal point (point an equivalent distance from metatarsal 2/3 marker along a vector normal to the foot plane formed by the heel, metatarsal 1 marker projected to height of metatarsal 5 marker, and metatarsal 5 marker, as the distance from the segment origin to the same foot plane) |
|  | Z-Axis | Metatarsal 2/3 marker to segment distal point (see y-axis) |
|  | Parent, DOF | Shank, 3 DOF |

Marker-based and markerless motion capture data were analyzed separately in Visual3D. Analog signals (force data) from the instrumented treadmill were imported into the markerless MoCap clothing and Sport clothing .c3d files using the Visual3D ‘Import_Signals_From_C3D_File’ pipeline command, allowing these signals to be used with the markerless kinematics to obtain joint kinetics and create force-based gait events. Analog signals were filtered using a low-pass critically-damped filter with a cut-off frequency of 20 Hz to match the kinematic filter cut-off frequency^9^, and the Visual3D ‘FP Auto Baseline’ pipeline command was used to clean the force platform signals. A vertical ground reaction force threshold of 50 N was used to detect foot contact events.

Kinematic and kinetic signals including joint positions, segment angles, joint angles, and internal net joint moments were calculated in Visual3D, and were time-normalized to 101 data points, representing one complete running gait cycle (0-100%) from heel strike to ipsilateral heel strike. Segment angles were calculated using a XYZ Cardan sequence relative to the global coordinate system and were included since they capture the 3D orientation of individual body segments. Joint angles were calculated using an XYZ Cardan sequence for all joints except the ankles, for which a XZY Cardan sequence was used. Joint angles were included since they capture the relative orientation of adjacent segments and are kinematic measures of interest for most users of motion capture. Joint moments were normalized to participant mass and resolved in the proximal segment. All data were analyzed further using python (version 3.10.6).

Four participants’ markerless Sport clothing trial videos had an inconsistent number of frames between cameras, preventing their processing in Theia3D; these participants were excluded from the kinematic comparisons, which then included 26 participants (12f/14m, mean (SD) age: 23.5 (4.0) years, height: 1.8 (0.1) m, mass: 69.9 (9.9) kg, BMI: 22.8 (2.9) kg/m^2^). Six participants’ center-of-pressure data from the instrumented treadmill were found to be erroneous, one of which was also part of the previously described kinematic exclusion group. The kinetic comparison sample therefore included 21 participants (10f/11m, mean (SD) age: 23.3 (3.8) years, height: 1.8 (0.1) m, mass: 69.9 (11.0) kg, BMI: 22.8 (2.8) kg/m^2^). Running gait cycles with marker tracking issues were excluded. For comparison between the markerless and marker-based motion capture, only matched gait cycles were used.

Differences between signals from corresponding running gait cycles for the concurrent marker-based and markerless MoCap clothing datasets were calculated and summarized across all participants as root-mean-square differences (RMSD). Participants’ mean signals for the asynchronous markerless MoCap clothing and Sport clothing conditions were used to assess differences between markerless clothing conditions and were summarized across all participants as RMSDs. The inter-cycle variability of the signals from each dataset was calculated^10^, and the inter-condition variability of markerless signals was also calculated to examine the reliability of markerless measurements across different clothing conditions (similar to ‘inter-session’ variability^10^).

### 2.3 Alternative Marker-Based Modelling

To investigate the effect of marker-based model definitions on kinematic and kinetic comparisons, results were calculated for two additional models. The first alternative model was our previous standard marker-based model that was used in prior publications**^3^** (Supplementary Materials, Table A). The second alternative model used the same set of markers but applied atypical segment definitions so that the resulting kinematics best aligned with the markerless results (Supplementary Materials, Table B). The modifications made to arrive at this model definition were selected on the basis of improving the agreement between the marker-based and markerless signals, and serve a purely illustrative purpose. This model was created by modifying the primary model used in the analysis, using techniques such as virtual markers that are offset relative to their real counterparts to define segment axes.

## 3. Results

The average self-selected running speeds for the participant samples used in the kinematic and kinetic comparisons were 2.42 (0.27) m/s and 2.39 (0.27) respectively.

### 3.1 Kinematics

Joint center positions from the concurrent marker-based and markerless MoCap clothing data had mean differences of less than 4 cm throughout the running gait cycle. The mean (SD) RMSD across all subjects were 2.12 (0.34) cm, 2.15 (0.63) cm, and 2.96 (0.87) cm for the ankle, knee, and hip joints, respectively (Figure 2). The markerless joint centers were positioned laterally of the marker-based joint centers for the ankle, knee, and hip, and distal relative to the marker-based joint centers for the ankle and hip. The sign of the anterior-posterior component was dependent on the phase of the running gait cycle and was distributed in both directions. The markerless ankle and hip joint centers were distal of their marker-based counterparts, while the knee was proximal during early swing phase and slightly distal during late swing and early stance.

**Figure 2:**
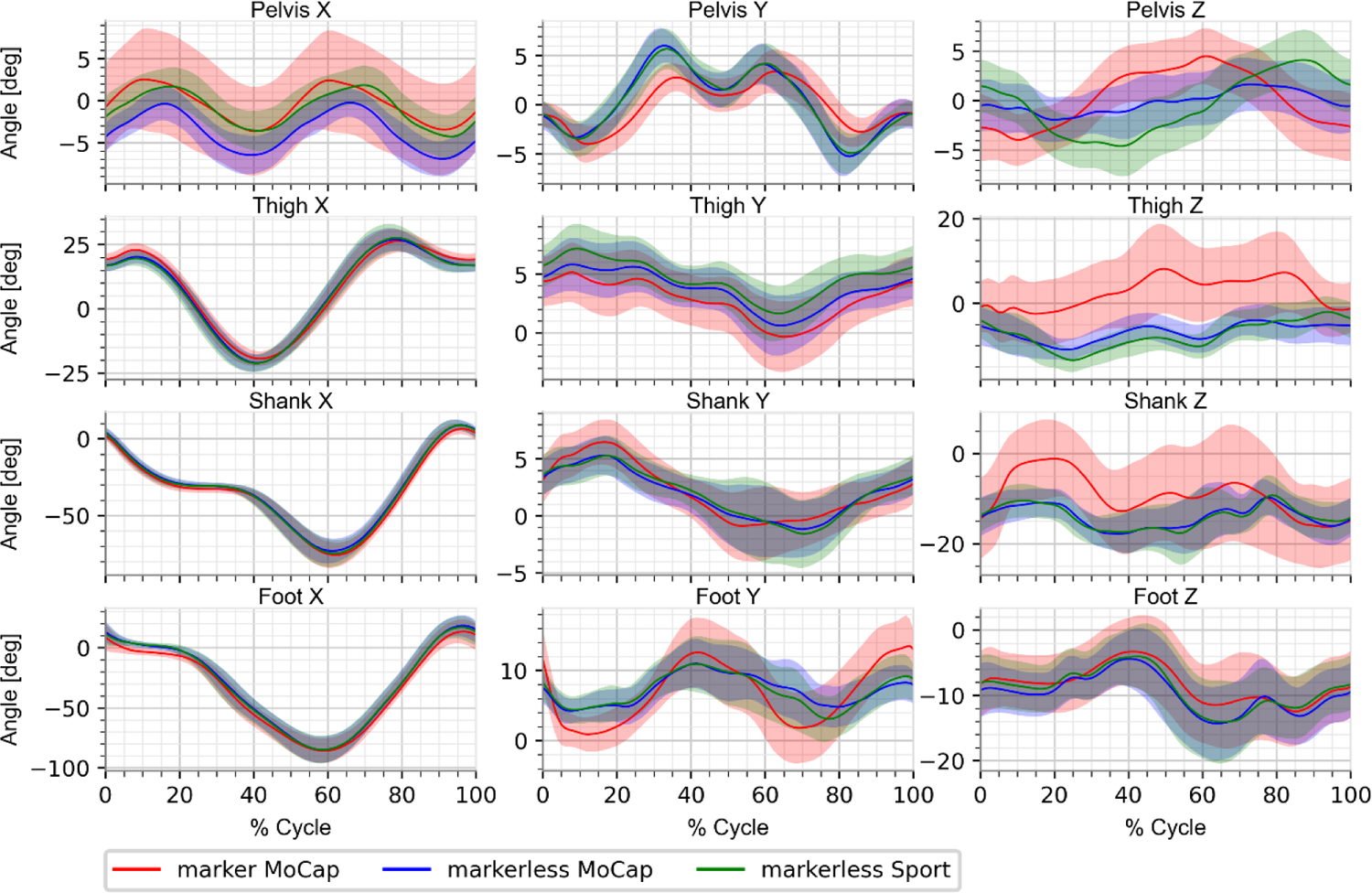
Mean right segment angles from marker-based motion capture (red), markerless motion capture under MoCap clothing condition (blue; concurrent with marker-based), and markerless motion capture under Sport clothing condition (green; asynchronous with marker-based and markerless MoCap clothing). Segment angle x-components (left column) represent rotations about the global coordinate system x-axis, z-components (right column) represent rotations about the segment local coordinate system z-axis, and y-components (middle column) represent rotations about an axis normal to the global x-axis and segment z-axis.

Lower limb segment angle x-components capture the largest magnitude of segment rotation during running, and showed very high visual similarity for the thigh, shank, and foot segments (Figure 3). Marker-based pelvis segment angle x-components showed greater variability than those from either markerless clothing condition. Segment angle y-component patterns were visually very similar for the pelvis, thigh, and shank segments, with larger differences between systems than between markerless conditions. The foot segment angle z-component patterns were also visually similar between all three datasets. The thigh and shank z-components, which capture rotations about the segment longitudinal axes, showed the greatest differences between motion capture systems. The pelvis segment angle z-components showed the greatest differences between the two markerless clothing conditions (MoCap, Sport) of any segment angle.

**Figure 4:**
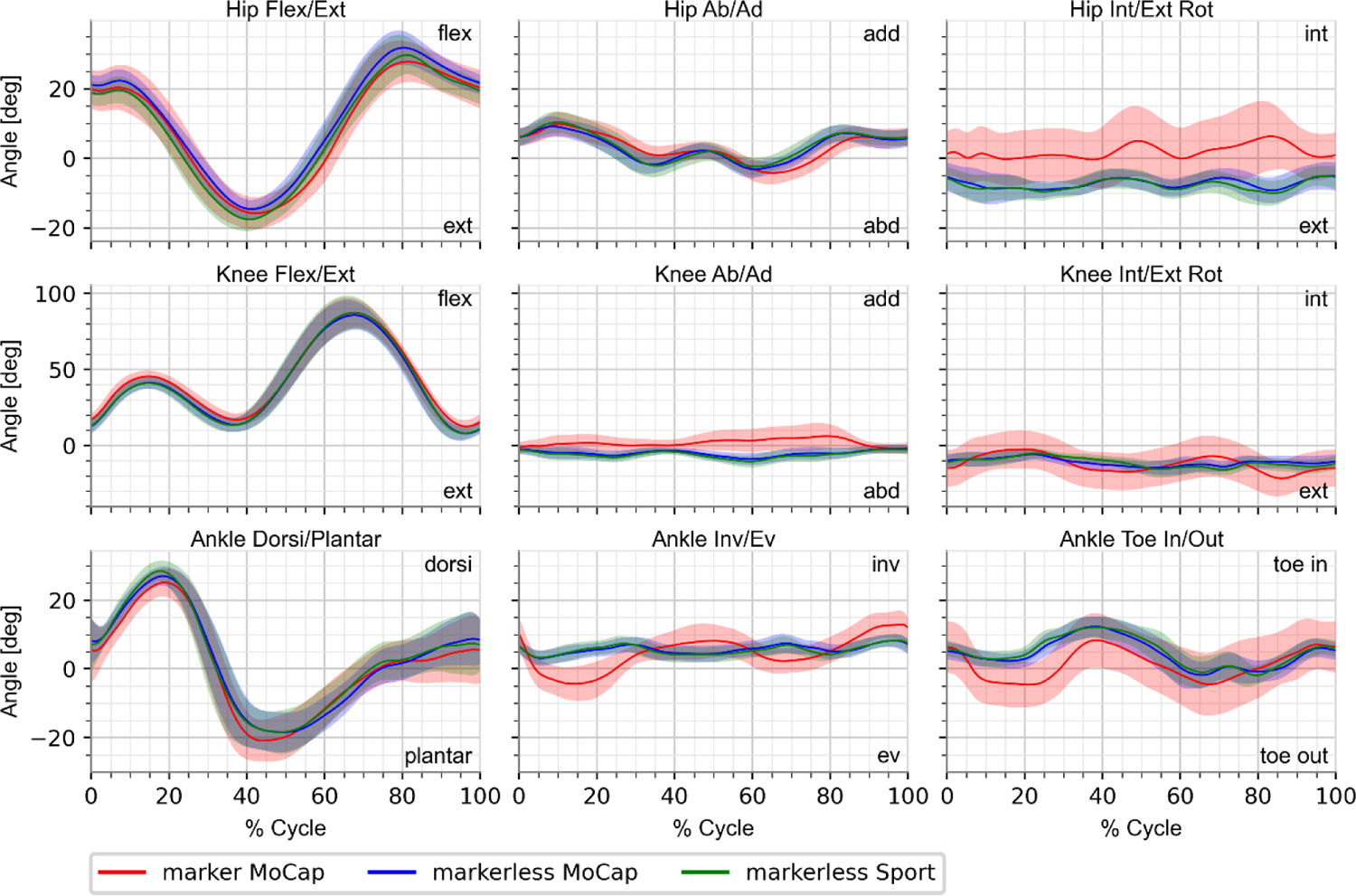
Mean right lower limb joint angles from marker-based motion capture (red), markerless motion capture under MoCap clothing condition (blue; concurrent with marker-based), and markerless motion capture under Sport clothing condition (green; asynchronous with marker-based and markerless MoCap clothing) using XYZ Cardan sequences for the knee and hip and a XZY Cardan sequence for the ankle.

Lower limb sagittal plane joint angles and hip ab/adduction (Figure 4) were visually similar across all three datasets, and had mean RMSDs of 5.2° or lower (Table 2). Ankle toe-in/toe-out patterns were also somewhat visually consistent between all three datasets during the swing phase, and had a mean RMSD of 8.0°. The hip internal/external rotation, knee ab/adduction and internal/external rotation, and ankle inversion/eversion angles captured different patterns and levels of variability between motion capture systems but were consistent between both markerless clothing conditions (mean RMSDs: 5-10°). The mean RMSD values for the asynchronous markerless data (MoCap clothing, Sport clothing) were all smaller than those for the concurrent marker-based and markerless data, indicating that the differences between markerless joint angle measures from different trials with different participant clothing were smaller than the differences between joint angles from marker-based and markerless systems from the same running gait cycles.

**Figure 4:**
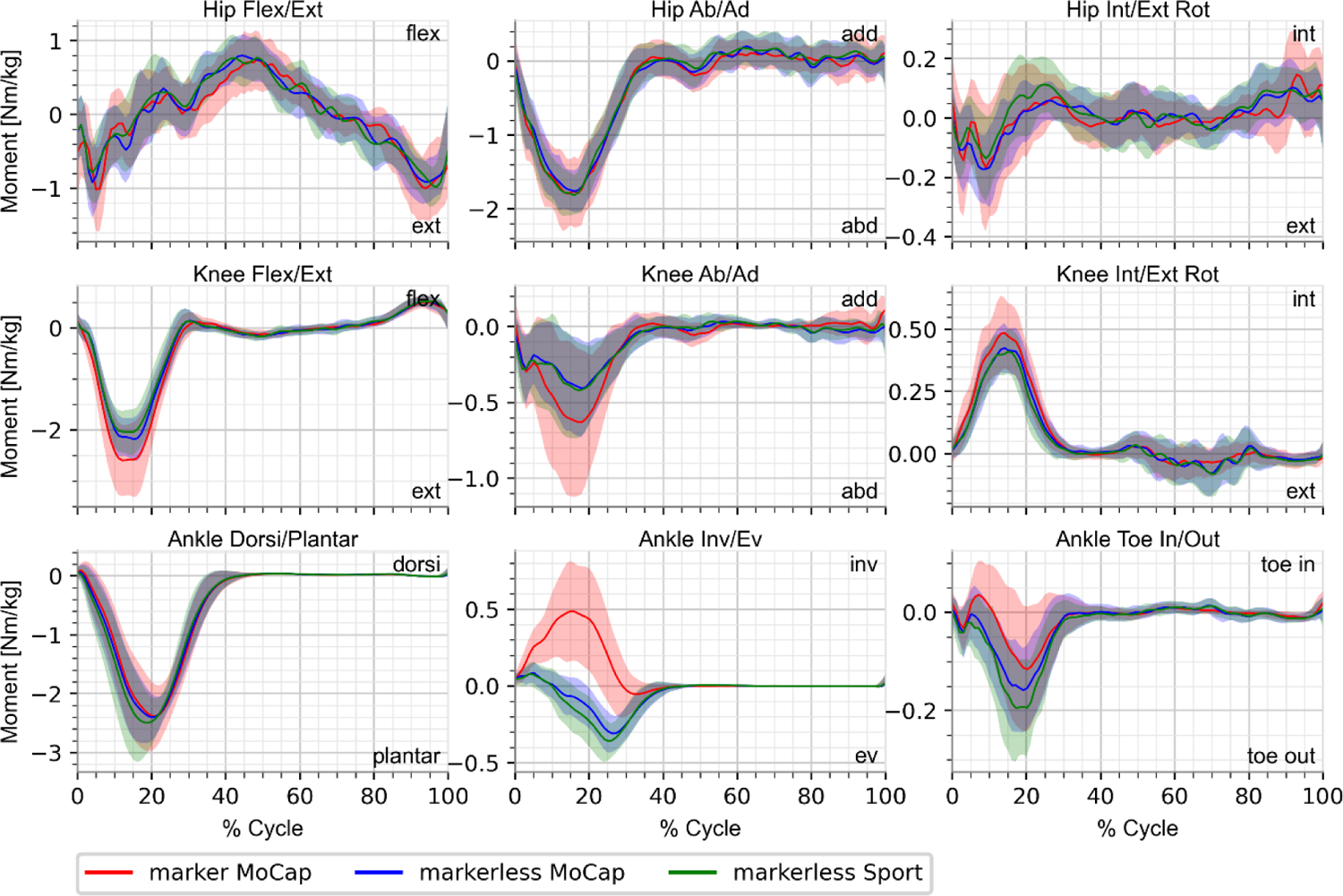
Mean right lower limb joint angles from marker-based motion capture (red), markerless motion capture under MoCap clothing condition (blue; concurrent with marker-based), and markerless motion capture under Sport clothing condition (green; asynchronous with marker-based and markerless MoCap clothing) using XYZ Cardan sequences for the knee and hip and a XZY Cardan sequence for the ankle.

**Table 2:** Mean lower limb joint angle and moment RMS differences across all participants, for the concurrent data comparison (markerless MoCap signals - marker-based MoCap signals) and the asynchronous comparison (markerless MoCap clothing signals - markerless Sport clothing signals).

|  |  | Ankle |  |  | Knee |  |  | Hip |  |  |
| --- | --- | --- | --- | --- | --- | --- | --- | --- | --- | --- |
|  |  | Fl/<br>Ex | Inv/<br>Ev | Toe-In/-<br>Out | Fl/<br>Ex | Ab/<br>Ad | Int/<br>Ext<br>Rot | Fl/<br>Ex | Ab/<br>Ad | Int/<br>Ext<br>Rot |
| Joint Angle<br>Differences<br>[deg] | Concurrent Data<br>Mean RMSD | 3.8 | 6.0 | 8.0 | 4.2 | 7.2 | 9.4 | 5.2 | 2.7 | 9.8 |
|  | Asynchronous<br>Data Mean<br>RMSD | 3.1 | 1.6 | 2.5 | 3.2 | 1.8 | 3.2 | 3.9 | 1.7 | 2.5 |
| Joint<br>Moment<br>Differences<br>[Nm/kg] | Concurrent Data<br>Mean RMSD | 0.13 | 0.25 | 0.03 | 0.24 | 0.12 | 0.06 | 0.29 | 0.22 | 0.09 |
|  | Asynchronous<br>Data Mean<br>RMSD | 0.13 | 0.05 | 0.03 | 0.13 | 0.08 | 0.05 | 0.25 | 0.18 | 0.07 |

Inter-cycle joint angle variability values were within 0.6° across all three datasets for all joint angles (Table 3). The average inter-cycle variability across all joint angles was slightly smaller for the marker-based data (1.3°) compared to those for both markerless datasets (1.5°). The inter-condition variability for joint angles between markerless conditions ranged from 1.3°-2.8° (mean 2.0°), leading to variability ratios of 1.2-1.7 (mean 1.3).

**Table 3:** Joint angle variability measures for each dataset and comparison, including: inter-cycle variability for marker-based MoCap, markerless MoCap clothing, markerless Sport clothing; inter-condition variability for all markerless data; and the variability ratio of markerless inter-condition to mean markerless inter-cycle variability.

|  |  | Ankle |  |  | Knee |  |  | Hip |  |  | Avg |
| --- | --- | --- | --- | --- | --- | --- | --- | --- | --- | --- | --- |
|  |  | Fl/<br>Ex | Inv/<br>Ev | Toe<br>-In/<br>-Out | Fl/<br>Ex | Ab/<br>Ad | Int/<br>Ext<br>Rot | Fl/<br>Ex | Ab/<br>Ad | Int/<br>Ext<br>Rot |  |
| Inter-Cycle<br>Variability [deg] | Marker-based<br>(MoCap clothing) | 1.5 | 1.2 | 1.4 | 2.1 | 0.8 | 1.3 | 1.3 | 0.9 | 1.3 | 1.3 |
|  | Markerless<br>(MoCap clothing) | 1.6 | 1.2 | 1.8 | 2.2 | 1.0 | 1.8 | 1.4 | 0.9 | 1.3 | 1.5 |
|  | Markerless<br>(Sport clothing) | 1.8 | 1.2 | 1.8 | 2.3 | 1.0 | 1.9 | 1.5 | 1.0 | 1.3 | 1.5 |
| Inter-Condition<br>Variability [deg] | Markerless<br>(MoCap, Sport<br>clothing<br>conditions) | 2.3 | 1.4 | 2.2 | 2.8 | 1.4 | 2.4 | 2.4 | 1.3 | 1.8 | 2.0 |
| Variability Ratio | Markerless Inter-Condition/Mean<br>Inter-Cycle | 1.4 | 1.2 | 1.2 | 1.2 | 1.3 | 1.3 | 1.7 | 1.3 | 1.4 | 1.3 |

### 3.2 Kinetics

Lower limb joint moments from all three datasets captured visually similar patterns for most joint moments, with differences observed in the knee flexion/extension and ab/adduction moments, ankle toe-in/toe-out moments, and large differences observed for the ankle inversion/eversion moments (Figure 5). The marker-based knee moments had greater peak magnitudes and greater variability compared to both markerless conditions in all three planes. The ankle inversion/eversion moments measured by the marker-based system were inversion moments across all subjects and had considerably greater variability compared to those measured by the markerless system, which were primarily eversion moments across all subjects in both conditions. Additionally, the ankle toe-in/toe-out joint moments measured by the marker-based system had a lower peak magnitude compared to both markerless conditions. The markerless MoCap clothing and Sport clothing conditions showed the biggest differences in the hip internal/external rotation moments, ankle inversion/eversion moments, and ankle toe-in/toe-out moments, for all of which the peak magnitude from the Sport clothing condition was greater than that of the MoCap clothing condition.

### 3.3 Alternative Marker-Based Modelling Results

The results of this study when two alternative marker-based models were applied are included in the Supplementary Materials (Tables A, B; Figures A-F). The results from the model used in previous studies^3^ were consistent with the results of that study, with slight variations due to the differences in task (walking versus running) (Supplementary Materials, Figures A, B, C). The results from the modified version of the model used here showed the same or greater similarity between marker-based and markerless systems for all segment angles, joint angles, and joint moments (Supplementary Materials, Figures D, E, F).

## 4. Discussion

The purpose of this study was to compare kinematic and kinetic measures of treadmill running from concurrent marker-based and markerless motion capture, and between asynchronous markerless motion capture under varying clothing conditions. Overall, we found comparable results between the simultaneous marker-based and markerless motion capture data and between clothing conditions for the markerless system.

There are several limitations of this study. Challenges with data collection resulted in some lost participants, but without a significant change in the demographics of the study population. The self-selected treadmill running speed of this group is lower than typical over-ground running speeds, which is likely due to previously observed perception differences in treadmill and over-ground running, and is consistent with previous findings^11,12^. Likewise, the framerate of 85 Hz for data collection is lower than is usually used in running studies but was a limitation of the hardware to allow maximum image resolution and synchronization across both motion capture systems. Although this is a relatively lower frame rate than is typically used to capture running, the purpose was to compare motion capture modalities rather than to study running biomechanics. Using different markerless motion capture hardware or capturing lower resolution videos would allow higher frequency data to be collected. We would anticipate that a higher running speed recorded at a higher framerate would have comparable results to those presented here. The order of the two clothing conditions was divided evenly across all participants, but not randomized and not considered in the data analysis. However, this would have no impact on the comparison of simultaneous marker-based and markerless data, and the lack of differences observed between clothing conditions implies that this was not an issue. Furthermore, the precise segment coordinate system definitions used by Theia3D are not publicly available, so our marker-based definitions that are intended to mimic the markerless segment definitions could possibly be improved upon; but in any case, the anatomical landmarks used by the markerless system are also not available and likely differ slightly in position from equivalent marker positions by nature of how they are created and tracked.

Joint position differences between marker-based and markerless concurrent data were found to be 3 cm or smaller for all joints, which is consistent with previous comparisons of these systems during treadmill walking^3^. Joint position component differences were also mostly consistent with those of Tang et al., however they did not report 3D Euclidean distances^6^.

Segment and joint angles demonstrated similar agreement between motion capture systems as in our previous study for treadmill walking gait^3^, and excellent agreement across asynchronous markerless clothing conditions. Sagittal plane joint angles between motion capture systems had mean RMSD of 5.2° or less for all three joints, consistent with the findings of Tang et al.^6^. The differences between systems measured as mean RMSD over the entire gait cycle are comparable to reported minimal detectable change (MDC) values for marker-based motion capture of treadmill running at initial contact^13^. Compared to the marker-based MDC values, the RMSD values between systems are 3.1° smaller (for ankle dorsiflexion) to 4.7° greater (for knee ab/adduction). The between-system agreement observed here is also consistent with previous comparisons of these systems during treadmill walking^3^, which showed larger differences for segment angles about the segment longitudinal axes, and for frontal and transverse plane angles. Kinematic and kinetic signals were highly consistent between the markerless MoCap clothing and Sport clothing conditions, except for pelvis segment angle x- and z-components. Interpreting kinematic differences between the two markerless motion capture clothing conditions is made challenging by the fact that they may result from actual changes in movement patterns between the two running trials or due to differences in markerless keypoint detections as a result of the change in clothing. The former was indeed the case for at least one participant, whose markerless joint angles were considerably different between the MoCap and Sport clothing conditions, and were confirmed from raw video data to have visibly greater ranges of motion in hip, knee, and ankle flexion for the Sport clothing condition (Supplementary Material, Figure H).

The RMSD values between clothing conditions, which could be considered analogous to a participant returning on a second day in different clothing, were smaller than published MDC values at initial contact^13^, with the exception of hip flexion (0.2° greater). These findings are consistent with a running repeatability study over three repeated visits at varying running speeds using the same markerless motion capture software^14^. Their findings and ours support the use of markerless motion capture methods to overcome the limitations of marker placement variability with marker-based motion capture, which has been a challenge for longitudinal running studies^15^.

Joint angle inter-cycle variability was small and similar across the marker-based data (1.3°) and both markerless datasets (1.5°). Inter-cycle variability is made up of natural stride-to-stride variation in subject biomechanics, as well as measurement variability inherent to the motion capture systems. Inter-condition variability across markerless clothing conditions was larger than the inter-cycle variabilities, and presented an average increase of 30% in measurement variability. The inter-condition variability is made up of inter-cycle variability as described above, plus natural inter-trial variability, measurement variability due to the varying participant clothing, and measurement variability inherent to the motion capture system. The inter-condition variability was smaller than the mean RMSD between concurrent motion capture systems for all joint angles, indicating that differences between markerless clothing conditions remained smaller than differences between motion capture systems.

Joint moments showed similarity in pattern and magnitude between systems and across clothing conditions for the ankle, knee, and hip sagittal plane moments, hip ab/adduction moment, and hip and knee internal/external rotation moments, while similar patterns of different magnitude were observed for knee ab/adduction and ankle toe-in/toe-out moments. However, the ankle inversion/eversion moments displayed different patterns, with the marker-based moments being mostly inversion moments while the markerless moments were mostly eversion moments. Differences in sagittal plane joint moment peak magnitudes between marker-based and markerless joint moments were mostly consistent with the findings of Tang et al., but they did not report frontal or transverse plane joint moments for either system^6^. The differences in ankle inversion/eversion and toe-in/toe-out moments between systems can be explained by differences in the foot and shank segment orientations, as the selection and orientation of the coordinate system in which joint moments are resolved can entirely change their direction^16^. For instance, when the ankle joint moments are resolved in the foot segment coordinate systems (Supplementary Material, Figure I) rather than the shanks (Figure 3), the marker-based inversion/eversion moments have a considerably reduced magnitude.

The results of this (and any) concurrent validation study of motion capture systems are highly dependent on the biomechanical models employed in the comparison. To investigate this, we applied two alternative models to the marker-based data that differed only in static segment definitions. Alternative model 1 (Supplementary Materials, Table A; Figures A-C) represented a standard unmodified marker-based model that has been previously published^3^. The model in Table 1 represents a slight compromise between standard marker-based and markerless segment definitions. Alternative model 2 (Supplementary Materials, Table B; Figures D-F) represents a model that was manually and intentionally modified to improve the agreement between the marker-based and markerless kinematic and kinetic signals. The modifications used in alternative model 2 were not extreme and remain within the realm of plausibility when considering the differences between marker sets and model definitions, and the variability of marker placement across operators. The results for this last model demonstrate the strong effects of segment definitions and marker placement on downstream biomechanical signals, as they demonstrate that a non-negligible portion of the differences in biomechanical signals can be attributed to local segment coordinate system misalignment and can be changed through relatively small model or landmark changes. The goal of this model was not to create or identify an optimally-modified model for matching marker-based and markerless signals, and it is entirely possible that different modifications could be implemented to greater success in matching the signals between systems. We advocate for the use of the automated coordinate system definition from Theia3D to facilitate standardization for multi-centre and longitudinal research, but found this a useful exercise to understand potential differences in how data has been collected from our historical marker-based approaches. This demonstrated the sensitivity of concurrent comparisons to these factors, which is an important point that should be understood when using and evaluating different motion capture solutions. These results highlight the challenges in combining data collected under different model definitions, and the potential benefit in using markerless motion capture to provide standardized results that can be pooled from multiple sites.

The results of this study indicate that Theia3D can measure treadmill running kinematics and kinetics with minimal effects from differing attire, and which are comparable to measures from marker-based motion capture. This study reiterates the influence of segment definition on resulting biomechanical signals. Careful consideration of these effects should be taken when conducting comparisons between measurement systems and relative to historical data that utilized potentially different segment definitions.

## 5. Acknowledgements

We thank Human Mobility Research Laboratory members with participant recruitment, data collection, and data processing. This research was supported by an Ontario Research Fund grant (ORF RE-04-047), Natural Sciences and Engineering Research Council of Canada (NSERC) Individual Discovery Grant, and NSERC Canadian Graduate Scholarship.

## Supplementary Materials

### 1. Effects of alternative model definitions for marker-based motion capture

#### 1.1 Alternative Model 1

**Table A:**
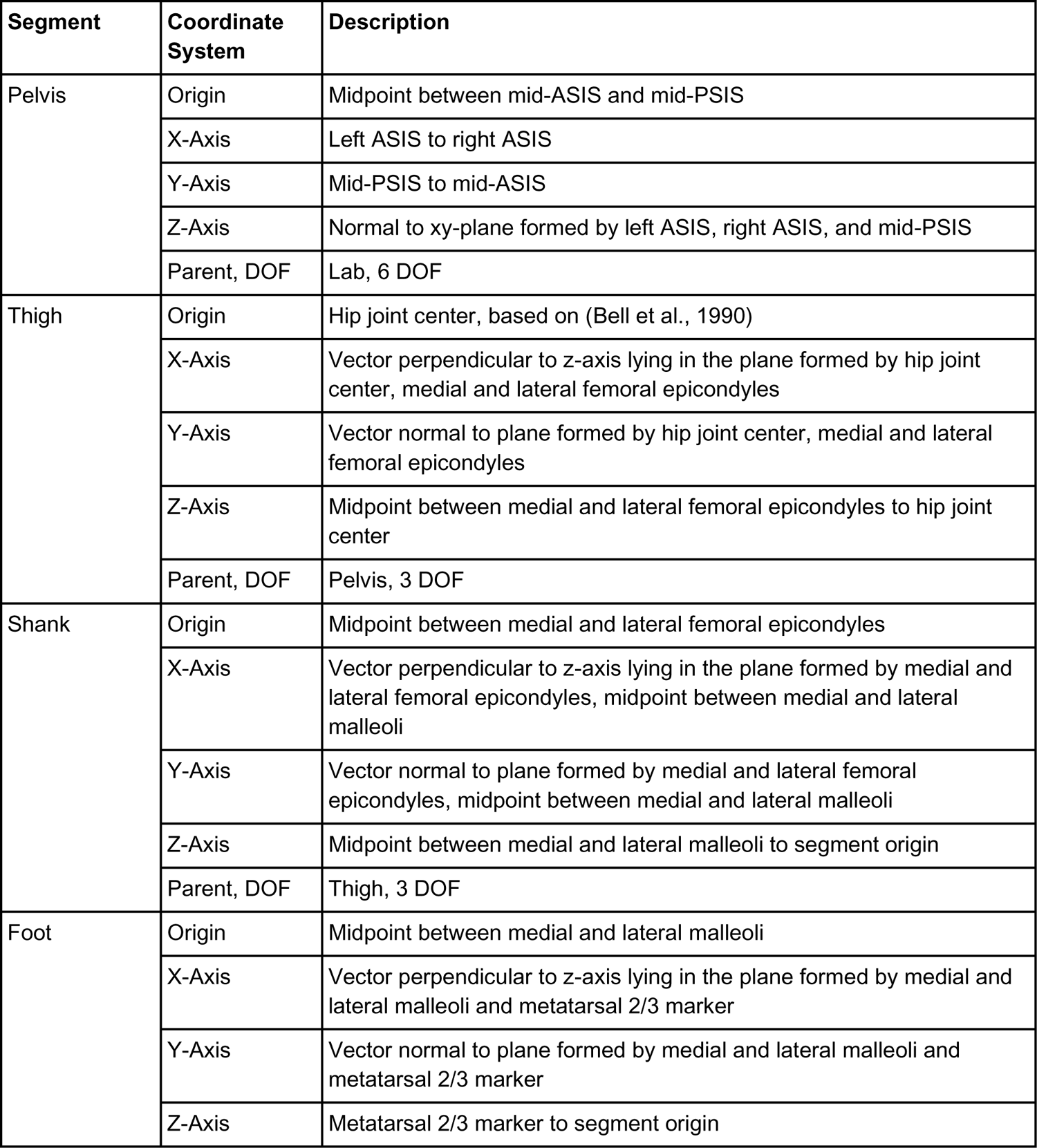
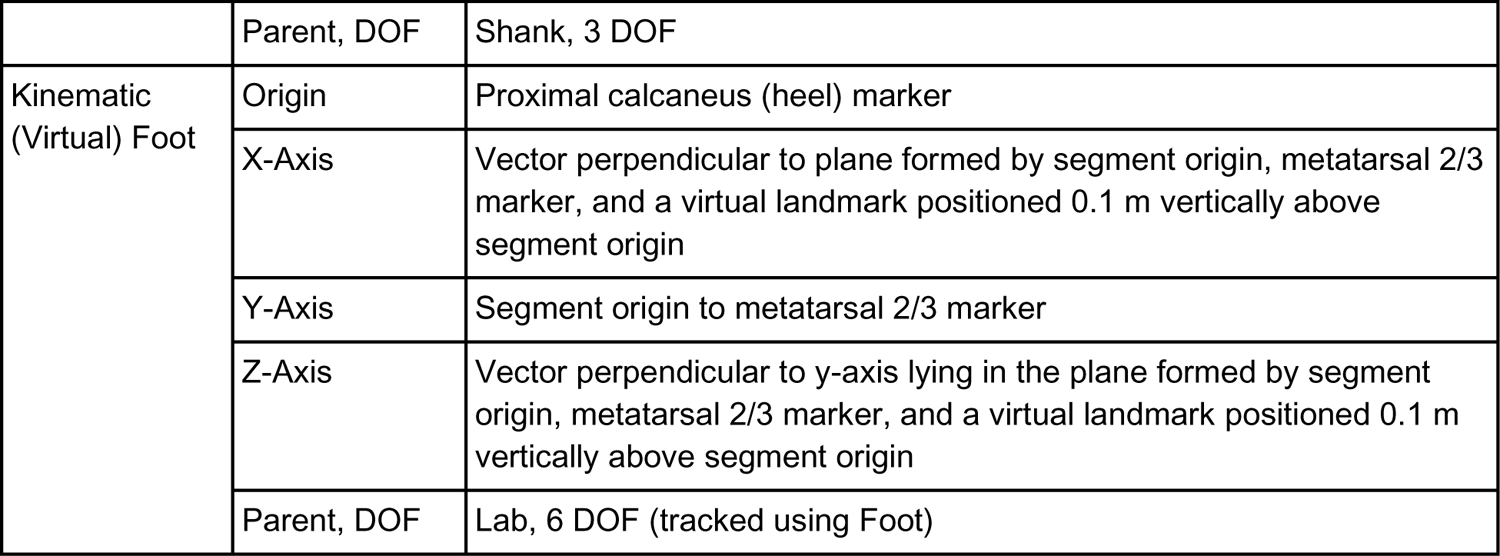
Marker-based model lower body segment coordinate system definitions and inverse kinematic model constraints for the model used in (Kanko et al., 2021a), corresponding to results shown in Figures A, B, C.

**Figure A:**
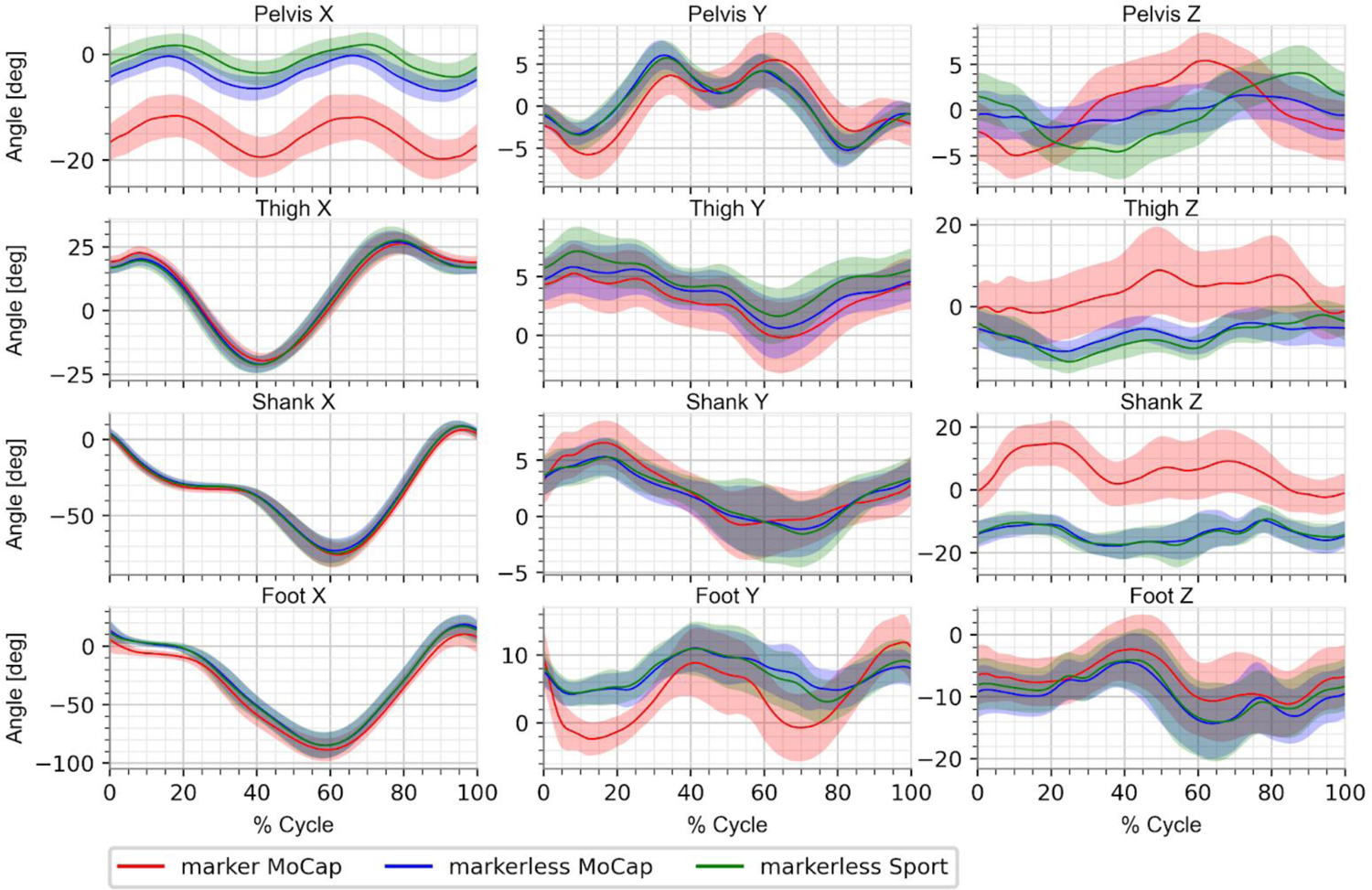
Mean lower limb right segment angles from marker-based motion capture (red), markerless motion capture under MoCap clothing condition (blue; concurrent with marker-based), and markerless motion capture under Sport clothing condition (green; asynchronous with marker-based and markerless MoCap clothing), using the marker-based model employed in (Kanko et al., 2021a).

**Figure B:**
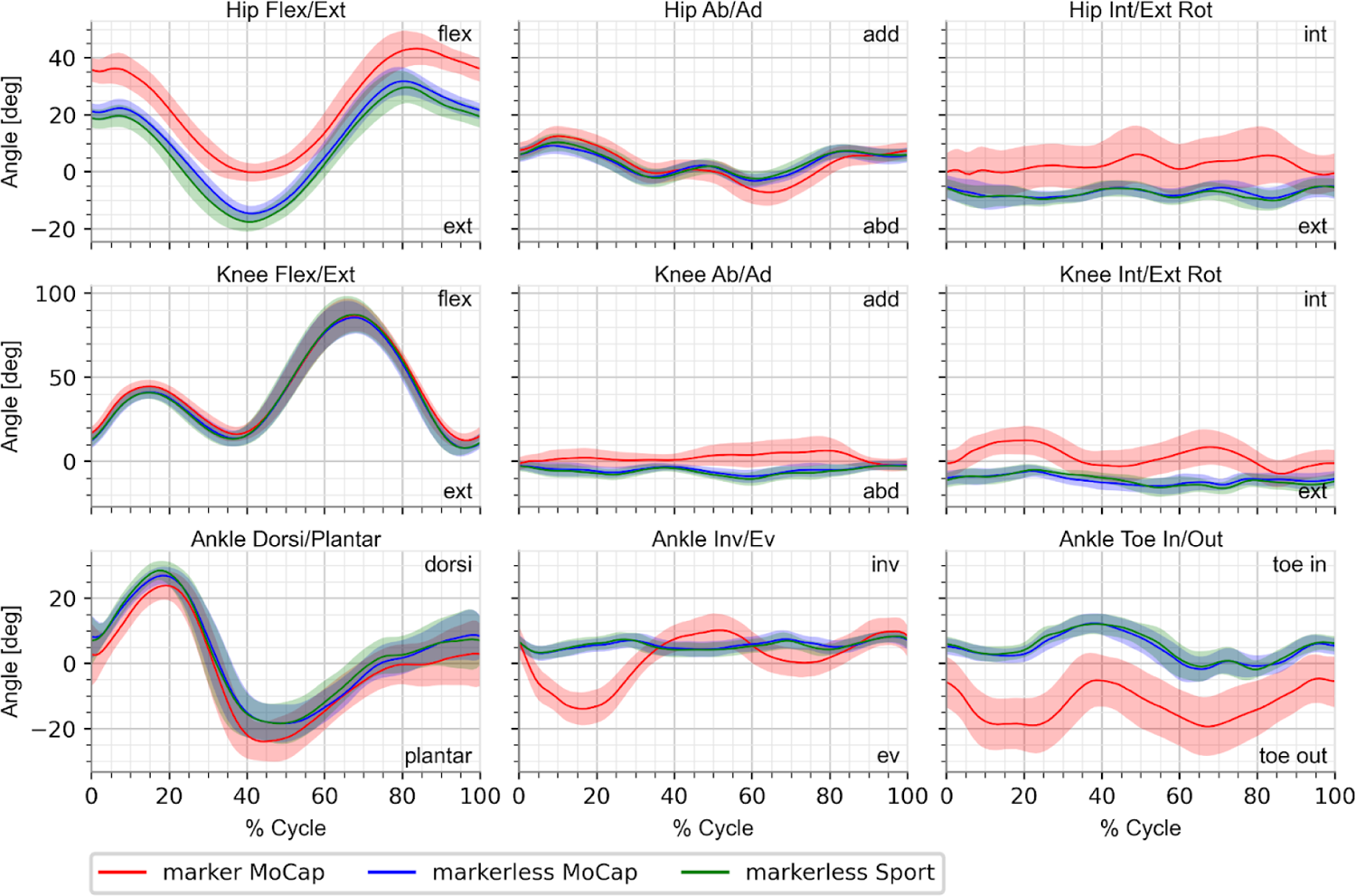
Mean lower limb right joint angles from marker-based motion capture (red), markerless motion capture under MoCap clothing condition (blue; concurrent with marker-based), and markerless motion capture under Sport clothing condition (green; asynchronous with marker-based and markerless MoCap clothing), using the marker-based model employed in (Kanko et al., 2021a).

**Figure C:**
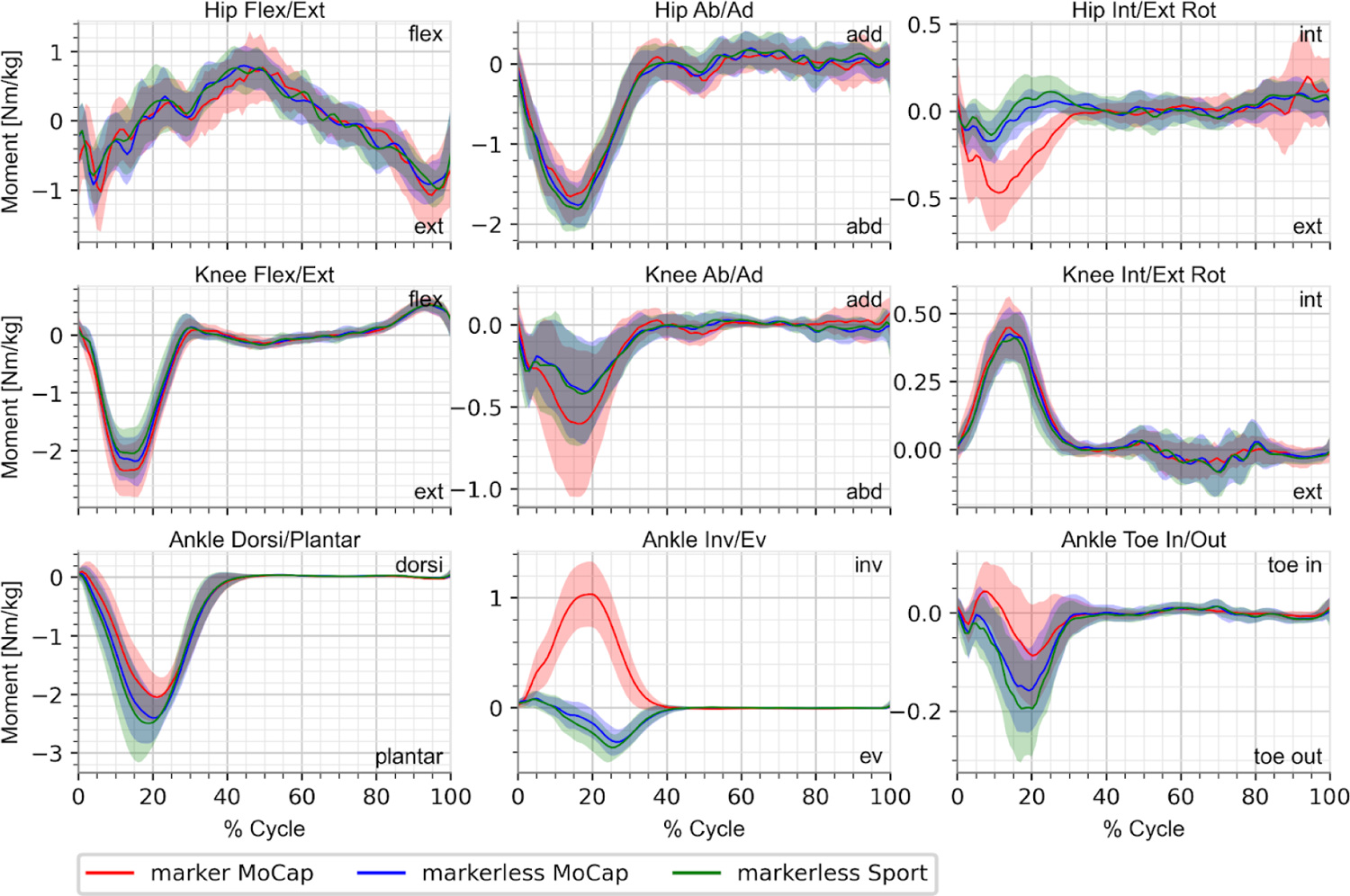
Mean lower limb right joint moments from marker-based motion capture (red), markerless motion capture under MoCap clothing condition (blue; concurrent with marker-based), and markerless motion capture under Sport clothing condition (green; asynchronous with marker-based and markerless MoCap clothing), using the marker-based model employed in (Kanko et al., 2021a).

#### 1.2 Alternative Model 2

**Table B:** Marker-based model lower body segment coordinate system definitions and inverse kinematic model constraints based on the model described in Table 1 determined through trial and error to better match the results from Theia3D for illustrative purposes, corresponding to results shown in Figures D, E, F.

| Segment | Coordinate System | Description |
| --- | --- | --- |
| Pelvis | Origin | Midpoint between mid-ASIS and mid-PSIS |
|  | X-Axis | Left ASIS to right ASIS |
|  | Y-Axis | Mid-PSIS to mid-ASIS |
|  | Z-Axis | Normal to xy-plane |
|  | Parent, DOF | Lab, 6 DOF |
| Kinematic (Virtual) Pelvis | Origin | Point at the height of midpoint between left and right iliac crest markers along vector between midpoint between left and right hip and midpoint between C7 and sternal notch markers. |
|  | X-Axis | Left hip joint center to right hip joint center |
|  | Y-Axis | Normal to xz-plane |
|  | Z-Axis | Segment origin to midpoint between C7 and sternal notch markers |
|  | Parent, DOF | Lab, 6 DOF (tracked using Pelvis) |
| Thigh | Origin | Hip joint center, based on (Bell et al., 1990) |
|  | X-Axis | Vector perpendicular to z-axis lying in the plane formed by segment origin, medial femoral epicondyle shifted 0.008 m anteriorly, and lateral femoral epicondyle shifted 0.008 m posteriorly |
|  | Y-Axis | Vector normal to plane formed by segment origin, medial femoral epicondyle shifted 0.008 m anteriorly, and lateral femoral epicondyle shifted 0.008 m posteriorly |
|  | Z-Axis | Midpoint between medial femoral epicondyle shifted 0.008 m anteriorly and lateral femoral epicondyle shifted 0.008 m posteriorly to segment origin |
|  | Parent, DOF | Pelvis, 3 DOF |
| Shank | Origin | Midpoint between medial femoral epicondyle shifted 0.02 m anteriorly and lateral femoral epicondyle shifted 0.02 m posteriorly |
|  | X-Axis | Vector perpendicular to z-axis lying in the plane formed by medial femoral epicondyle shifted 0.02 m anteriorly, lateral femoral epicondyle shifted 0.02 m posteriorly, and midpoint between medial and lateral malleoli |
|  | Y-Axis | Vector normal to plane formed by medial femoral epicondyle shifted 0.02 m anteriorly, lateral femoral epicondyle shifted 0.02 m posteriorly, and midpoint between medial and lateral malleoli |
|  | Z-Axis | Midpoint between medial and lateral malleoli to midpoint between medial femoral epicondyle shifted 0.02 m anteriorly and lateral femoral epicondyle shifted 0.02 m posteriorly |
|  | Parent, DOF | Thigh, 3 DOF |
| Foot | Origin | Midpoint between medial and lateral malleoli |
|  | X-Axis | Vector perpendicular to z-axis lying in the plane formed by segment origin, segment distal point (see y-axis), and a shifted lateral malleoli marker (shifted vertically by the height of the medial malleoli marker – 0.95* height of the lateral malleoli marker - 0.0186 m) |
|  | Y-Axis | Segment origin to segment distal point (point an equivalent distance from metatarsal 2/3 marker along a vector normal to the foot plane formed by the heel, metatarsal 1 marker projected to height of metatarsal 5 marker, and metatarsal 5 marker, as the distance from the segment origin to the same foot plane) |
|  | Z-Axis | Vector perpendicular to y-axis lying in the plane formed by segment origin, segment distal point (see y-axis), and a shifted lateral malleoli marker (shifted vertically by the height of the medial malleoli marker – 0.95* height of the lateral malleoli marker - 0.0186 m) |
|  | Parent, DOF | Shank, 3 DOF |

**Figure D:**
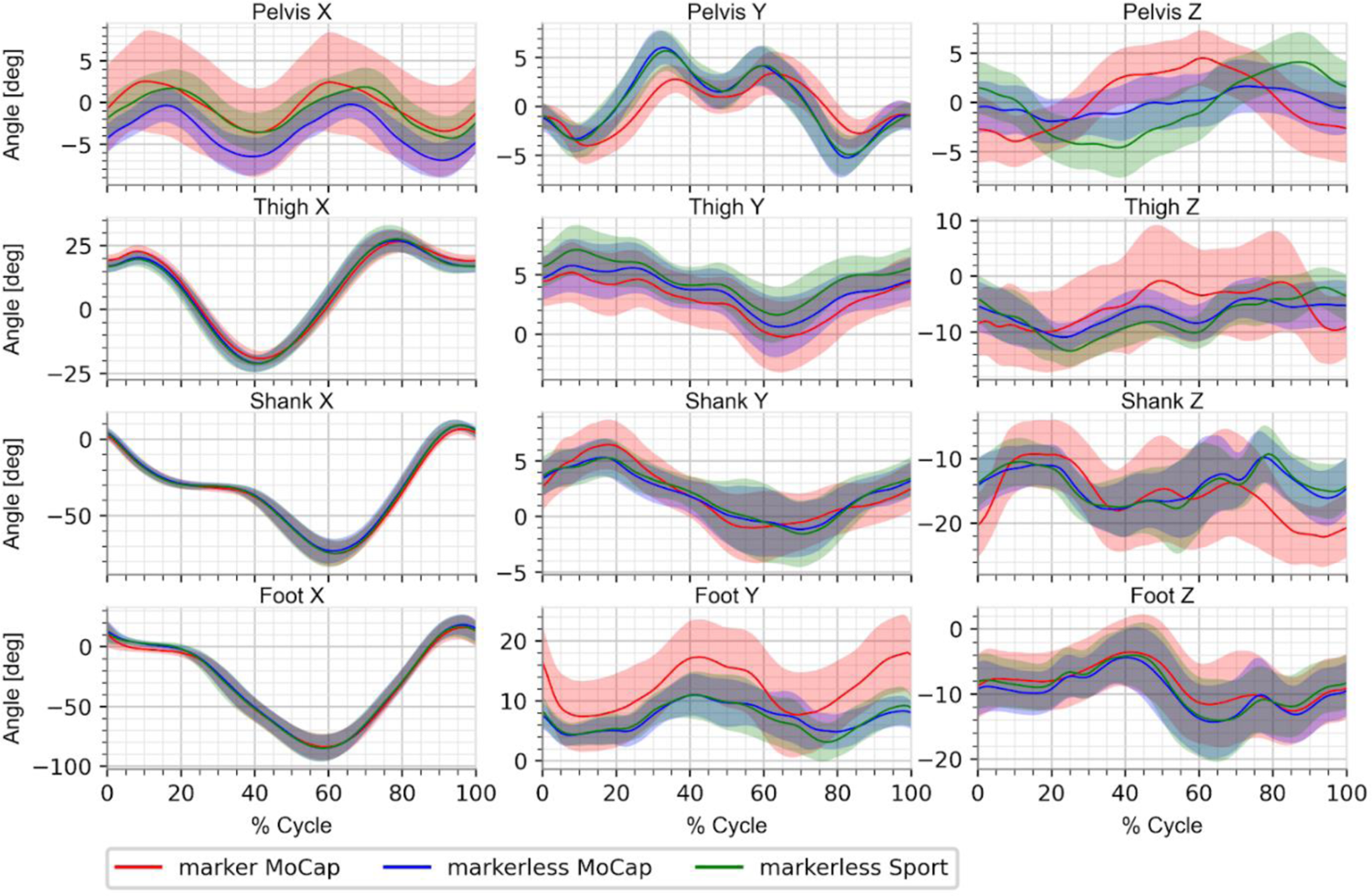
Mean lower limb right segment angles from marker-based motion capture (red), markerless motion capture under MoCap clothing condition (blue; concurrent with marker-based), and markerless motion capture under Sport clothing condition (green; asynchronous with marker-based and markerless MoCap clothing), using the marker-based model defined in Table B to better match the results from Theia3D for demonstration purposes.

**Figure E:**
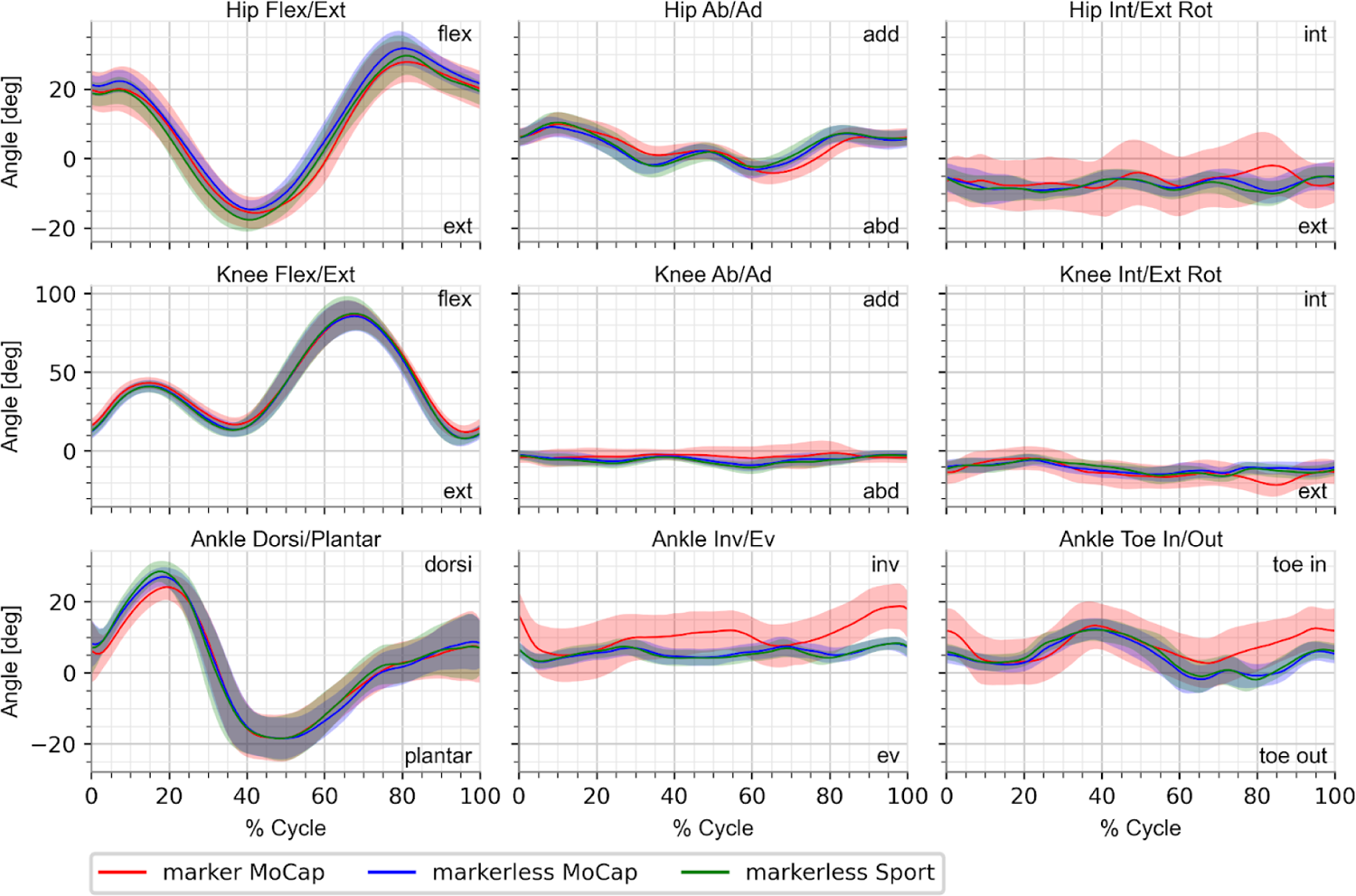
Mean lower limb right joint angles from marker-based motion capture (red), markerless motion capture under MoCap clothing condition (blue; concurrent with marker-based), and markerless motion capture under Sport clothing condition (green; asynchronous with marker-based and markerless MoCap clothing), using the marker-based model defined in Table B to better match the results from Theia3D for demonstration purposes.

**Figure F:**
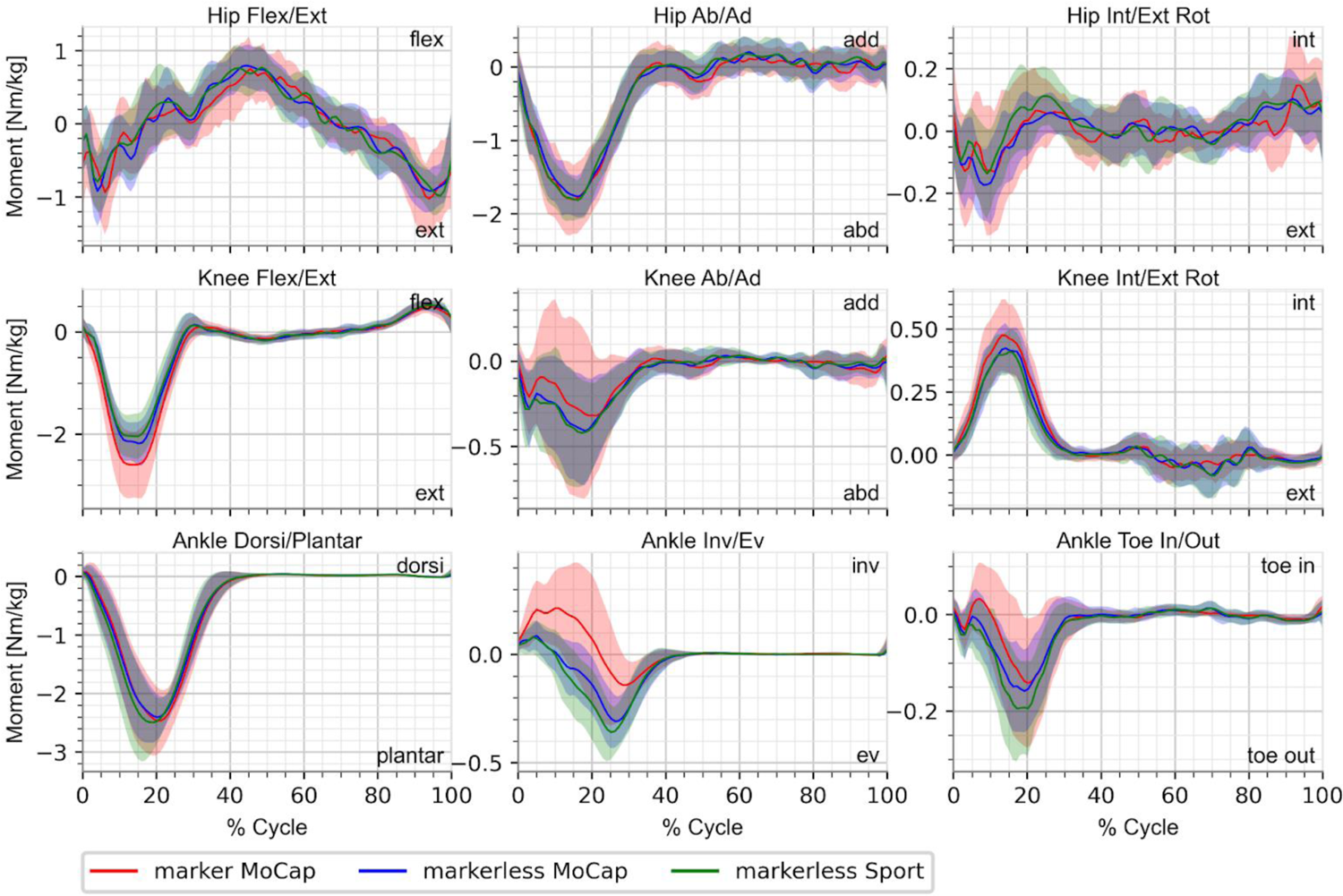
Mean lower limb right joint moments from marker-based motion capture (red), markerless motion capture under MoCap clothing condition (blue; concurrent with marker-based), and markerless motion capture under Sport clothing condition (green; asynchronous with marker-based and markerless MoCap clothing), using the marker-based model defined in Table B to better match the results from Theia3D for demonstration purposes.

#### 2. Example individual participant kinematic waveforms

Raw joint angle data for a representative participant with consistent running biomechanics (Figure G) and a participant with altered running biomechanics when wearing MoCap clothing compared to their self-selected Sport clothing (Figure H).

**Figure G:**
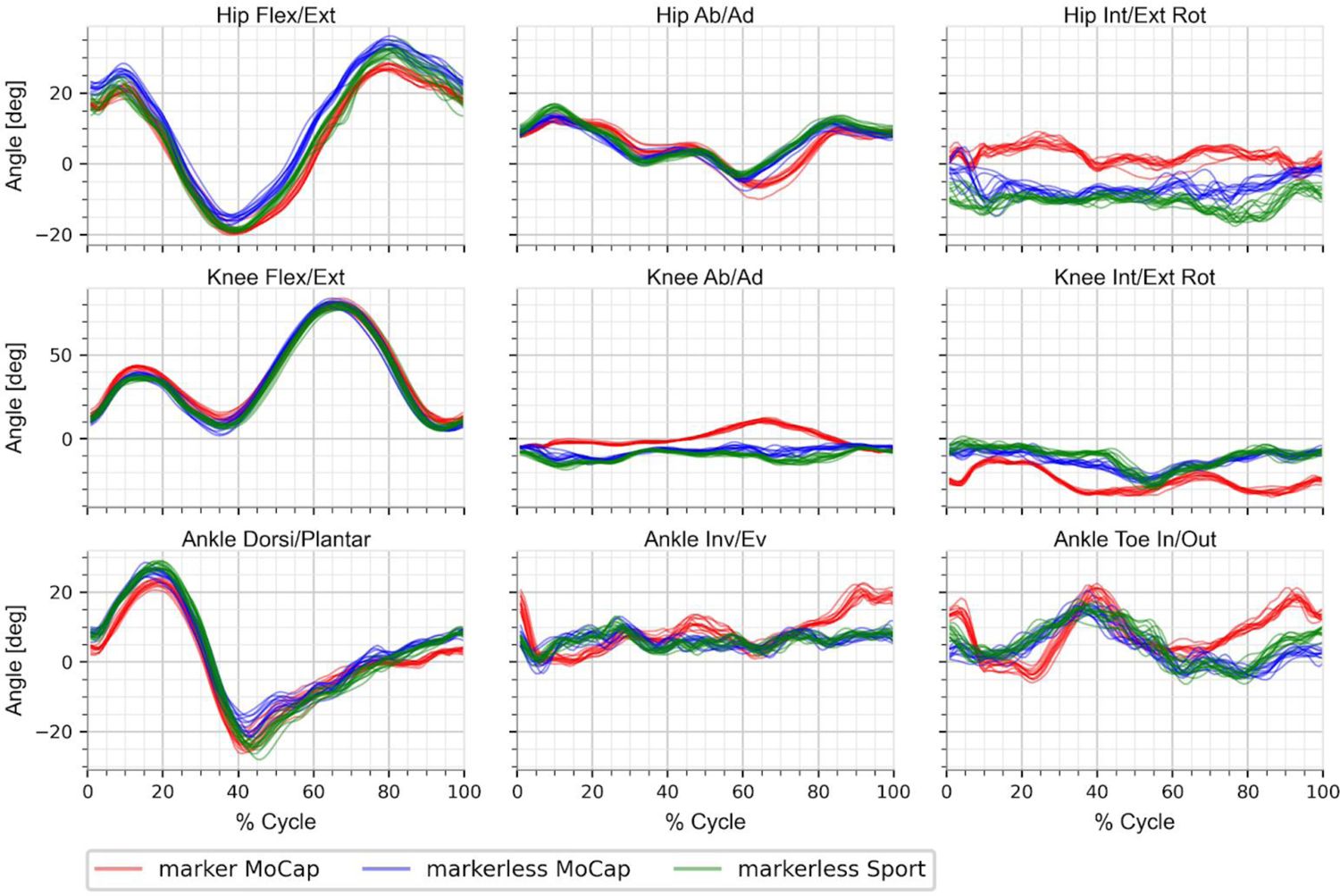
Raw lower limb right joint angles for one representative participant, from marker-based motion capture (red), markerless motion capture under MoCap clothing condition (blue; concurrent with marker-based), and markerless motion capture under Sport clothing condition (green; asynchronous with marker-based and markerless MoCap clothing).

**Figure H:**
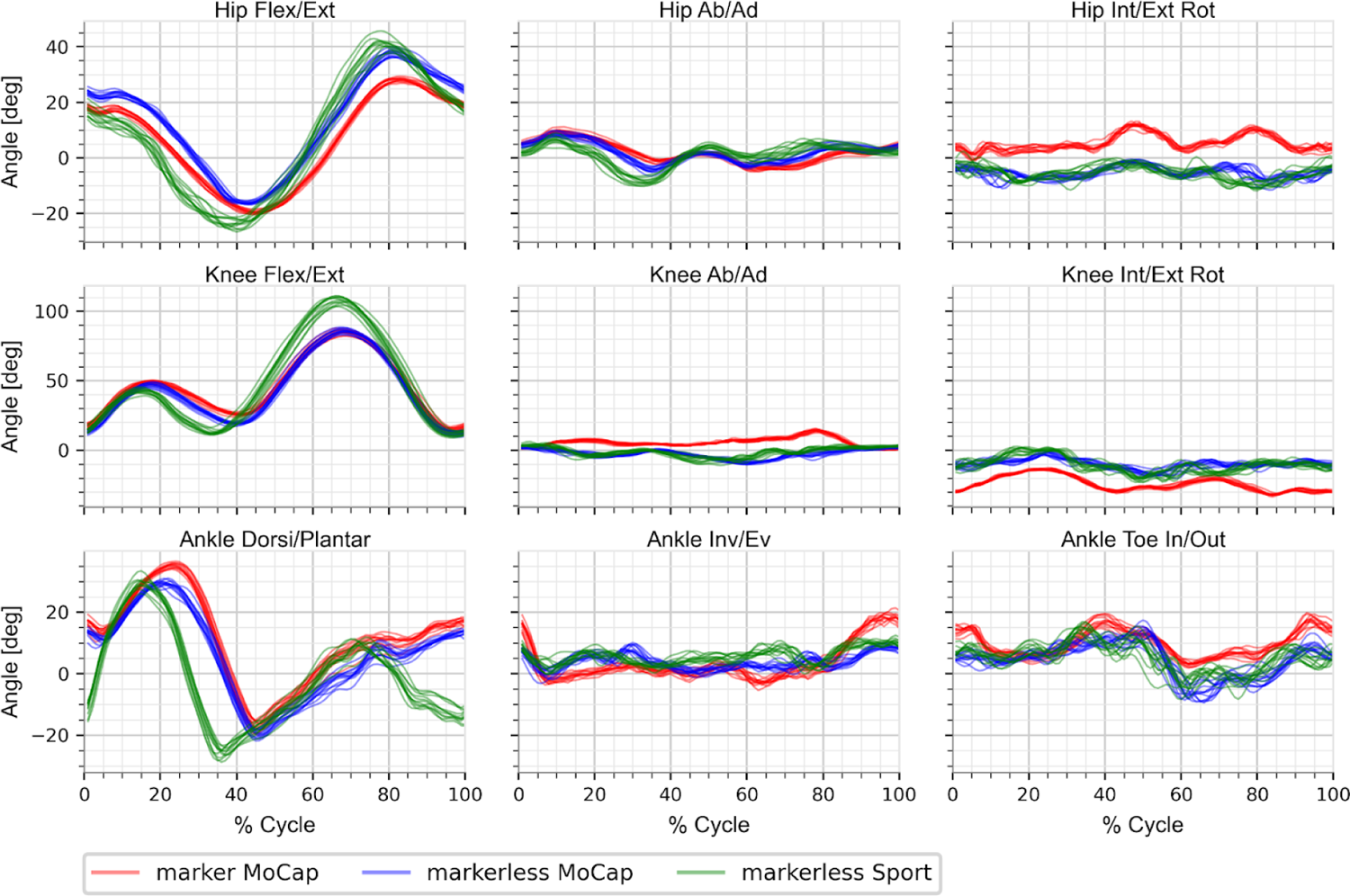
Raw lower limb right joint angles for one participant who used different biomechanical patterns between the MoCap and Sport clothing conditions, as visually confirmed from the raw markerless videos. Joint angles shown from marker-based motion capture (red), markerless motion capture under MoCap clothing condition (blue; concurrent with marker-based), and markerless motion capture under Sport clothing condition (green; asynchronous with marker-based and markerless MoCap clothing).

#### 3. Effect of ankle moments resolved in foot segment coordinate system

**Figure I:**
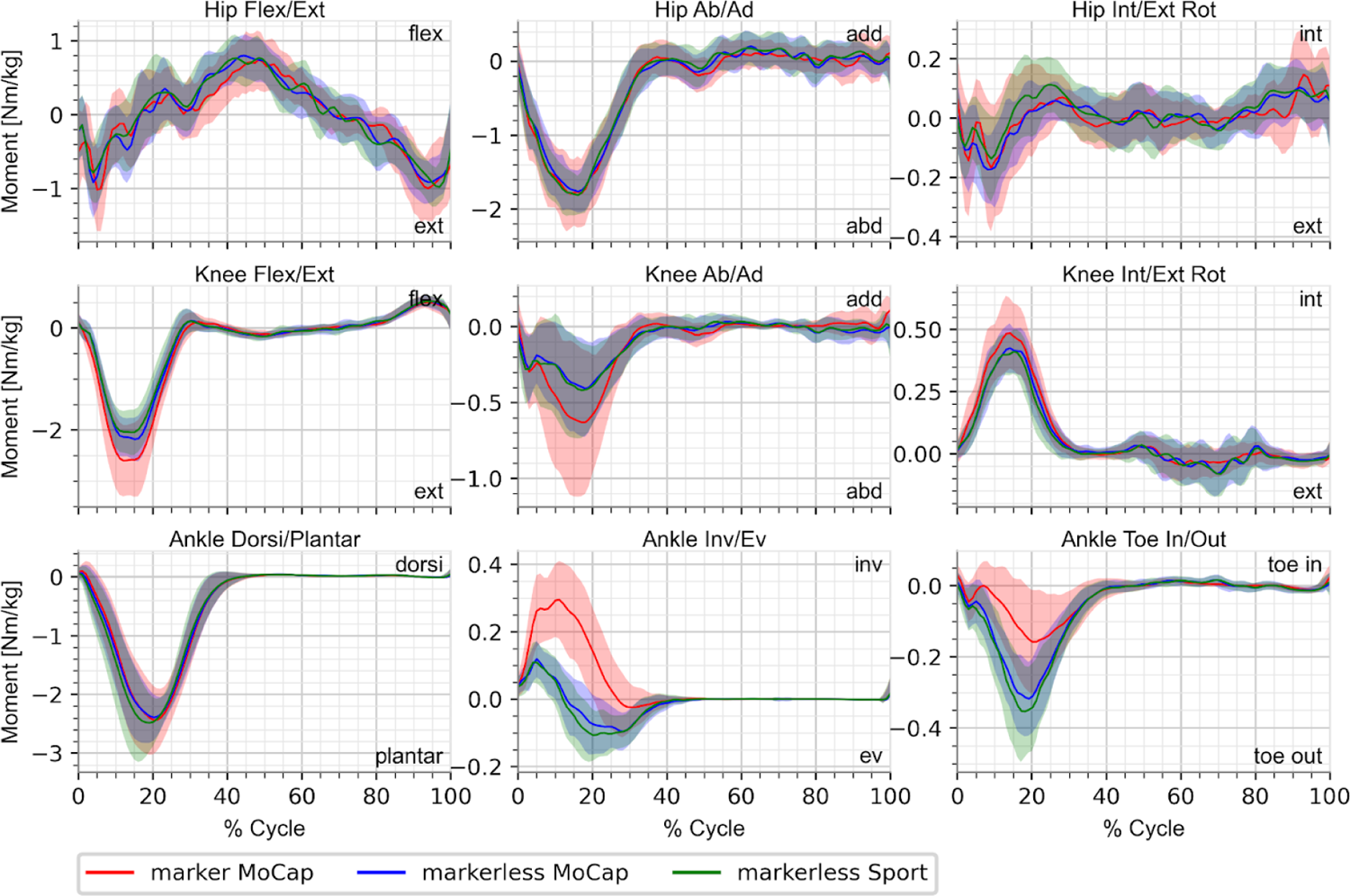
Mean lower limb right joint moments from marker-based motion capture (red), markerless motion capture under MoCap clothing condition (blue; concurrent with marker-based), and markerless motion capture under Sport clothing condition (green; asynchronous with marker-based and markerless MoCap clothing), with the ankle joint moments resolved in the foot segment coordinate system.

